# Introducing the technique of group-handling in C57BL/6NCrl mice using a novel device “Citizen- Chitra Mus-Mobile”, a cage-transfer cum enrichment system, reduces stress, anxiety and time taken for cage changes and enhances voluntary interaction

**DOI:** 10.1101/2025.05.31.656571

**Authors:** VS Harikrishnan, Arvind Kumar Prajapati, Kamalesh Mehta

**Affiliations:** Sree Chitra Tirunal Institute for Medical Sciences and Technology, Biomedical Technology Wing, Satelmond Palace, Poojappura, Trivandrum, Kerala, India; Citizen Industries, Navrangpura, Ahmedabad, Gujarat, India

**Author notes:** Corresponding author, Sree Chitra Tirunal Institute for Medical Sciences and Technology, Biomedical Technology Wing, Satelmond Palace, Poojappura, Trivandrum, Kerala, India.

**Keywords:** Group-handling, laboratory mice, stress, welfare, refinement

## Abstract

It has been proven that tail-lifting mice can cause stress, anxiety and aversive behavior, and this finding has made scientists develop less stressful techniques like cupping and tunnel-handling. All these techniques have a disadvantage in that they can only handle mice individually and not in groups. This study produces scientific evidence on the advantages of group handling of laboratory mice for the first time. Group handling using the novel device significantly reduced the time taken and number of attempts taken for cage changes, a major reason that prevented tunnel-handling from being adopted globally. Shifting mice in groups using a device named “Mus-Mobile” resulted in significantly lower fecal corticosteroid levels, higher voluntary interaction time and the open field activity and time spent in the central arena were not altered when compared with tail lifting and tunnel handling in C57BL/6NCrl mice. Body weights were also not different when Mus-Mobile was employed with respect to the other groups. This work shows that group-shifting is the next horizon for animal welfare, as this device can revolutionize the handling of laboratory mice to bring in better animal wellbeing.

## Introduction

Tail handing of laboratory mice can result in lesser anxiety, higher stress responses and low voluntary interaction with handlers (1) and this can even affect the outcome of research results including pharmacokinetic data (2). Mice handled by tail showed varying results in comparison with tunnel-handled ones in behavioral studies, as they reduce mice’s responses to reward and thereby exhibiting reduced pleasure-seeking behavior (3). The stress thus induced can affect reproducibility of studies in biomedical research (4). So, more depressive like state is observed in mice handled using tail-lifts and this can affect test data generated in pharmacology research (3). Animal welfare and scientific quality of data produced, both, are two important sides of biomedical research (2) and refinement in handling is an important area that shall be worked upon with to improve the quality of science and welfare. These findings eventually led to the discovery of techniques like handling mice using cupped hands and using tunnels (1). However, an international thematic survey revealed that, time constraints were cited by most respondents as a reason for not using tunnel handling to handle laboratory mice (4). Further, there exists no proven techniques to handle laboratory mice in groups and in turn, no data on its advantages on animal welfare.

On these grounds, a novel device was conceptualized, fabricated and tested to handle laboratory mice in groups. It was hypothesized that the same device can also serve as an enrichment device for the animals. After a habituation period of 7 days, in a 7-day study, It was decided to investigate the fecal corticosteroid concentration, time taken to perform cage changes, voluntary interaction, and activity in the open field when the novel device was employed and to compare it with tail lifting and tunnel handling. The novel device was named as “Citizen Chitra Mus-Mobile” as it handles the mice in groups and was applied for Indian Patent (Indian Patent Application No. 202241061704 dated 29/10/2022 entitled “Mice transfer box tunnel system with securing gates) and design registration was also done (New Design registration application No. 457573-001; dated 02.05.2025). While refinement is made, care shall also be exercised to avoid any unwanted effects in research results or functionality of the animal model (5) and so fluctuations in body weights was also assessed as part of the study. The study aimed to establish that group handling using the novel device can improve animal welfare by reducing stress, speeding up the process of cage changing and thereby refine the existing techniques in mice handling with scientific data.

## Materials and Methods

### Ethical approval, animals and their care

Permissions for animal experimentation was obtained from the Institutional Animal Ethics Committee of Sree Chitra Tirunal Institute for Medical Sciences and Technology (SCTIMST) and the work was entitled “Testing the acceptance of a novel mice transfer and enrichment box tunnel system with securing gates and its effect on stress levels and normal behaviour of C57BL/6J, BALB/c and Swiss Albino mouse colonies” with a Form B Sanction Number SCT/IAEC/536 /FEB/121/2025 dated 07th February 2025. SCTIMST’s animal facility is registered with the Committee for the Control and Supervision of Experiments on Animals (CCSEA), a statutory body under the legal frame work of Indian animal experimentation rules. C57BL/6NCrl mice of six weeks of age with both the sexes in equal proportion from Charles River Laboratories that was sourced through Hylasco Bio, India Limited, Bangalore was used for the work. The animals were housed in a barrier-maintained facility with controlled environment (RH between 30 to 70 %; Temperature 22±2°C and 10-15 fresh air changes per hour). Health monitoring in the facility was performed as per FELASA guidelines (6) and the colony was healthy and free from any adventitious agents. The animals were housed in polysulfone IVC cages (Type C cages, Citizen Industries, Ahmedabad, India) of 800 sq. cm cage floor area and 18 cm cage height with 700 g of autoclaved corncob as bedding (Sparcobb, India) and were fed with *ad libitum* food (Safe rodent diet, D131, Augy, France) and water. Aspen wood-based nesting material (10 g net weight per cage) and a chewing wood block (KSPL Wood nesting material and KSPL S Brick for mice, KSPL OPC Ltd, Himachal Pradesh, India) each was supplied to all cages for promoting species specific behaviour. Cage changes were performed once in five days, and the lighting was limited to not exceed 325 Lux at 1 m height from the floor with an automated 14:10 D: L cycle. Noise levels were kept below 85dB. Three well trained and experienced personnel were involved in the handling and testing experiments who were equally assigned randomly to all the three groups. Handling and behavioral tests were performed between 10.00 am and 2.00 pm in all cases.

### Study Design

72 C57BL/6NCrl mice were completely randomized and blocked were equally assigned into three groups with equal number of males and females featuring per group (n= 24 per group with 12 males and 12 females). Sample size was decided based on results observed from previous behavioral studies. Further, the current study was also fixed based on an apriori sample size calculation. It was decided to observe the animals daily and in case of cage-mate aggression and resultant injuries, especially as seen in C57BL/6NCrl males, to exclude such mice from the study. The group I was assigned to be handled using conventional tail base grip (tail-lifting) and group II with tunnel handling (1). Group III was assigned to be handled in groups using the novel device as described in the next paragraph. A red standard polysulfone mouse igloo (Bio-Serv Mouse Igloo) with entry ports was kept in all the cages assigned as group I. A red standard mouse made up of polysulfone tunnel was kept permanently in the cages assigned as group II. The novel polysulfone Mus-Mobile was placed in all cages assigned as group III.

### The novel device for group-handling of mice – “the Citizen Chitra Mus Mobile” and its operation

The device features a two-part design (Part A and Part B, Fig. 1.) made in polysulfone with an optional and detachable two-gated design with interior tunnel and raised platform has outer dimensional measurements of 10 cm (L) x 10 cm (W) x 7.5 cm (H) making it compatible with all sizes of individually ventilated and conventional cages for laboratory mice. It includes circular entry ports that mimic burrows and a rectangular outer structure that maximizes space efficiency. To provide enrichment in terms of increased activity and hence to alleviate boredom, it has a climbing platform at the height of 3.5cm from the base with a ladder that serves as a dual-purpose play station (Fig.2.). The device incorporates multiple air holes for continuous airflow, ensuring a comfortable environment for the mice. Constructed from durable materials such as polysulfone or polyetherimide, the device is resistant to gnawing, autoclavable and designed for longevity. The front section (Part A) can be detached as shown in Fig. 1. for aiding effective cleaning and subsequent sterilization. The front and rear doors are fitted with multiple rings functioning as hinges for smooth opening and closing, enhancing operational efficiency during mice transfer. The doors are free to be opened by mice from inside the device and must be kept in an open position, by keeping it rest on the top when they are kept inside cages. The length of the doors is designed in such a way that, once the doors are kept open and when used in mice cages (up to 20 cm height) are closed using its top grill, these gates cannot close even accidentally when mice manipulate the doors. With rounded edges and polished interiors, the device ensures safety for animals and can accommodate up to 6 adult mice.

The device can be placed inside the mice cages permanently.

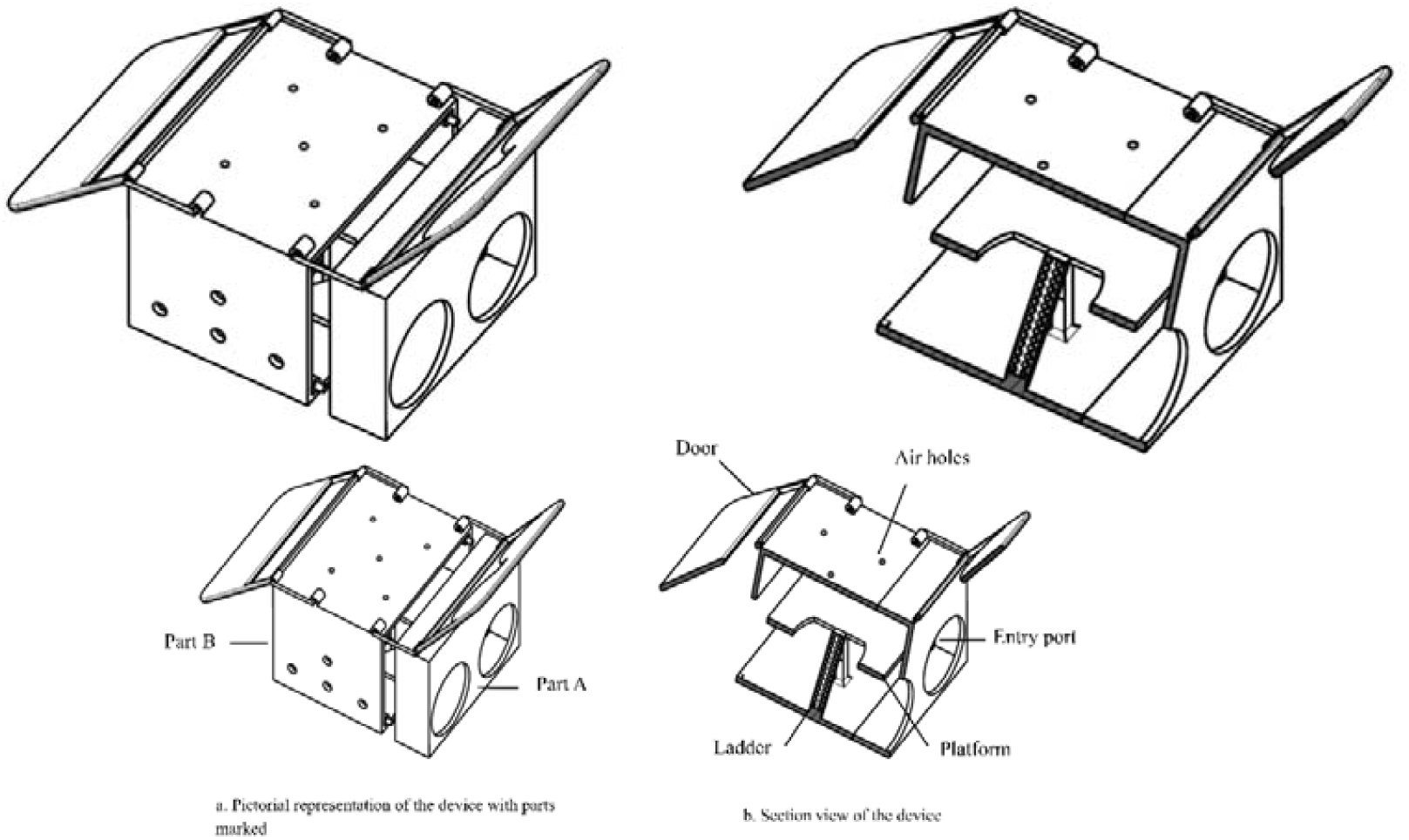
Fig 1a and 1b 1a Pictorial representation of the device 1b Section view of the device

Nesting material if supplied to mice, will be used for nesting in close proximity of the device (Fig.2a.). A minimum of 24 hours shall allow animals to get acclimatized to the device before which handling shall not be attempted. Once the handler opens up the cage, the mice will move out of the nest (Fig.2. b.). Once the handler attempts to handle the mice, most of the animals (75%) will voluntarily move fastly into the device to take shelter (Fig.2c.). The handler can close one side of the device using his palm of the non-dominant hand and direct the remaining ones to the device by carefully ushering and guiding the mice using the other hand to enter the device (Fig.2d.). Once all the animals have entered the shifting device, the other side also shall be closed using the palm and the animals can be lifted and shifted as a group (Fig.2e and f.).

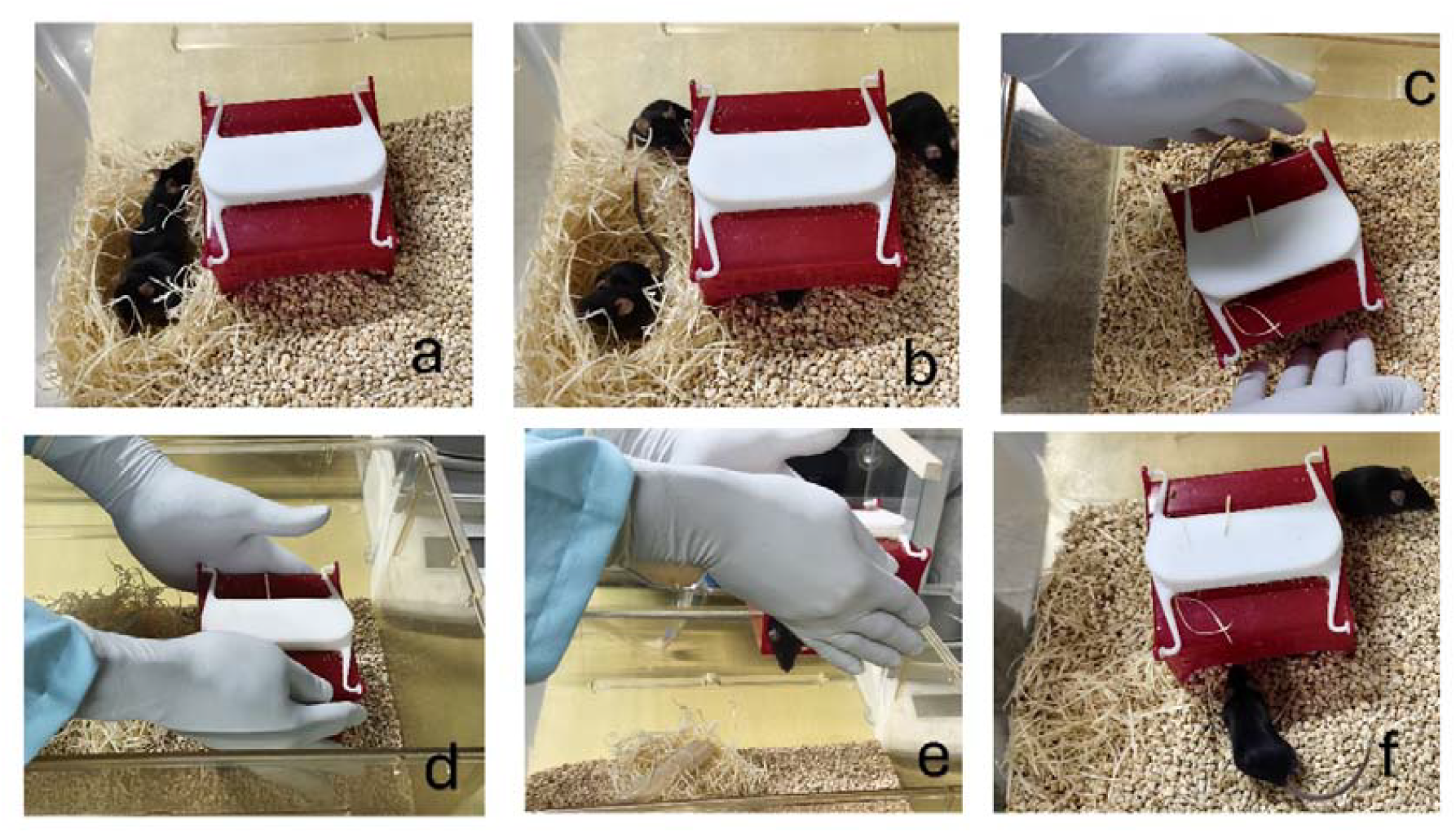
**Fig.2a**. The mice build nests in close vicinity to the novel device **Fig.2b**. The mice move out of the nest as soon as they notice the handler’s presence **Fig.2c**. Once the handler attempts to handle, 75% of the animals takes shelter inside the novel device **Fig.2d**. The handler can usher the remaining mouse/mice to enter the device and close its entry and exit ports with both his/her palms and pick the nesting box up **Fig.2e**. The group of mice can be shifted to the clean cage or to wherever the handler wants it to be **Fig.2f**. The mice get out of the Mus-mobile in its new cage

If in one attempt all the animals have not entered into the device and is left out in the first attempt to pick up the device for cage shift, the device with animals shall be placed in the new cage. The same device was not be attempted to be taken back by tapping on it to make the indwelling animals to move out, since this could cause stress. Our protocol stated that, in such rare occasions, a second spare device shall be kept ready to be used to collect the remaining animals. After the spare device is used to shift the remaining animals, the handler has to wait until the mice move out of it voluntarily to remove the second device from the cage to which animals get shifted. Furthermore, it is advised not separate Part-A from Part-B of the device or re-attach while animals are inside. This is to prevent the animals getting hurt with any of their body parts being stuck in between.

### Timeline of the study

The animals were received from the vendor at 4 weeks of age and after a quarantine of one week, they were acclimatized for 7 days. Until this point, all the animals were handled using tail-lifting. During this period, fecal pellets were collected in 3 days to obtain an average baseline corticosteroid level. Body weights were obtained in week 6 and the animals were randomized and blocked and were grouped into 3 groups and from here onwards, they were handled only using respective techniques for a week. After 24 hours after the animals were introduced to the novel device and tunnel, in week 6, cage changing time, number of attempts to change all animals from one cage to another and voluntary interaction was studied. Tests to estimate the cage changing time and number of attempts while changing cages were repeated after 24 hours to get adequate sample size and also to reinforce the trend of data obtained. Open field test was done 24 hours after the conduct of voluntary interaction test. This time gap was allocated to overcome any residual effects from previous tests. One week after being introduced to the assigned techniques, 24-hour fecal pellets were collected and were stored in -20°C for analysis of corticosteroid metabolites. Body weights were obtained in week 7 after all handling was completed during week 6.

### Tests done to assess efficiency of handling, stress, anxiety like behavior and welfare

#### Time taken for cage changing to assess efficiency of handling

The time taken to transfer the animals from one cage to another to assess efficiency of handling was performed. For this, the cage with the animals and the empty cage of same dimensions were kept side by side in their longitudinal axis touching each other and perfectly in line. The animal cage’s water bottle, top cover, and wire lids were completely removed and kept aside. The count started when the handler has completed the shifting of nesting material, handling device/tunnel/igloo and chewing block from the cage to the clean cage and when his hands start to move towards the device/tunnel to pick up the animals. In the case of tail lifting, when the handler makes the movement intended to pick up the first animal. The count ended when the shifting of all the 4 animals in the cage is completed. The time in seconds was noted and was compared between groups.

#### Number of attempts to move animals between cages to assess efficiency of handling

The number of attempts to move out all the four mice from one cage to the next was calculated for each technique and was compared between groups.

#### Voluntary Interaction Test

The test was performed as described before by Hurst and West (7). Time spent interacting with the device, tunnel or gloved hand closely was measured in seconds and was compared between groups.

#### Open field Activity

An open field arena with raised borders with dimensions 65 × 50 × 20 cm was used for the open field test. The inner area of 30×45 cm was assigned as central arena. The mice were let free at the center of the arena using their respective handling technique, one animal at a time. The entire open field was divided into 4 equal quadrants and the number of crossings between quadrants were counted and expressed as animal’s activity. The number of crossings between quadrants and the time spent in the central arena was recorded and compared between groups using a video camera fixed above the open field, and recording was done for 3 minutes. The fecal pellets of an animal were removed and the arena was cleaned using ethanol wipes before another animal was introduced into the field.

#### Faecal Corticosteroid assay

Faecal corticosteroid assay was performed using Cayman chemical corticosterone ELISA Kit, (Item number 501320, Cayman chemical, MI, USA). Average baseline value from three readings was taken and difference between the corticosteroid concentrations measured after handling using the assigned technique was found out and compared between groups.

#### Body weights

Body weights before and after handling by the designated methods were compared between groups.

### Statistical Analysis

Each individual animal constituted an experimental unit for all the tests except cage changing time, number of attempts and fecal corticosteroid metabolites where cage was the unit. Statistical analysis was performed using GraphPad Prism version 10.5.0 for Windows, GraphPad Software, Boston, Massachusetts USA, www.graphpad.com. Data is represented as individual animals representing each dot in the scatter plot in all cases except for cage changing time, number of attempts and corticosteroid assay where the dot represents the cage. Mean is shown with the horizontal bar and SD the whiskers. All the data was tested for normality using Kolmogorov-Smirnov test and normal data was tested using ANOVA and post-hoc multiple comparisons were done using Tukey’s test. Non-normal data was compared using Kruskal-Wallis test with Dunn’s test employed to carry out pairwise comparisons. P<0.05 was considered as a statistically significant difference between groups.

## Results

There were no cage-mate aggression and resultant exclusions from the study. All groups completed the observation period in a healthy and satisfactory manner.

### Body weights

Body weights did not differ between groups on day one (prior to being subject to handling procedures) or day seven (after handling procedures).

### Cage to cage changing time

**Fig 3.**
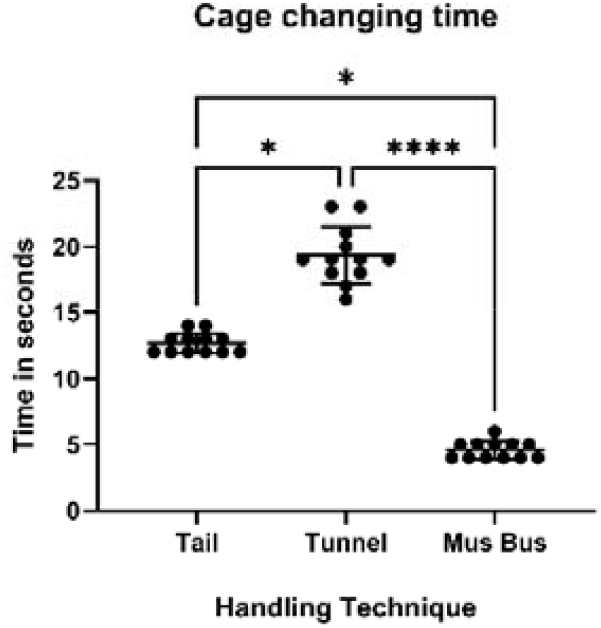
Mus-Mobile was significantly swift in transfer-handling of animals in groups when compared with tail and tunnel handled groups of mice (p<0.0001) (Kruskal Wallis statistic= 31.59).

### Number of Attempts

**Fig 4.**
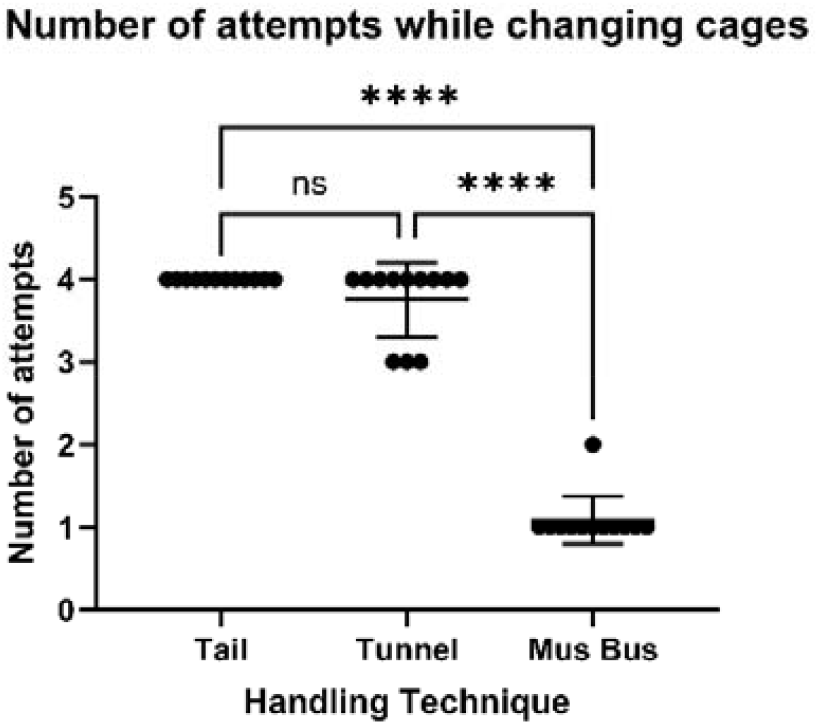
Mus-Mobile shifted all the 4 animals in one-go in most of the changes and was significantly advantageous when compared with the other two groups (P<0.0001) (Kruskal Wallis statistic= 30.84).

### Voluntary Interaction Test

**Fig 5.**
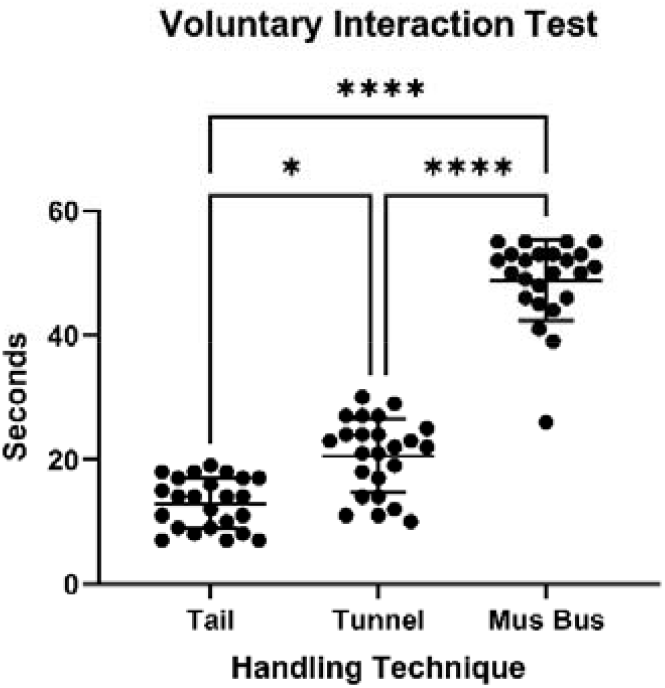
The animals interacted more voluntarily when Mus-Mobile was used for transfer when compared with the other two groups (P<0.0001) (Kruskal Wallis Statistic = 54.23).

**Fig 6.**
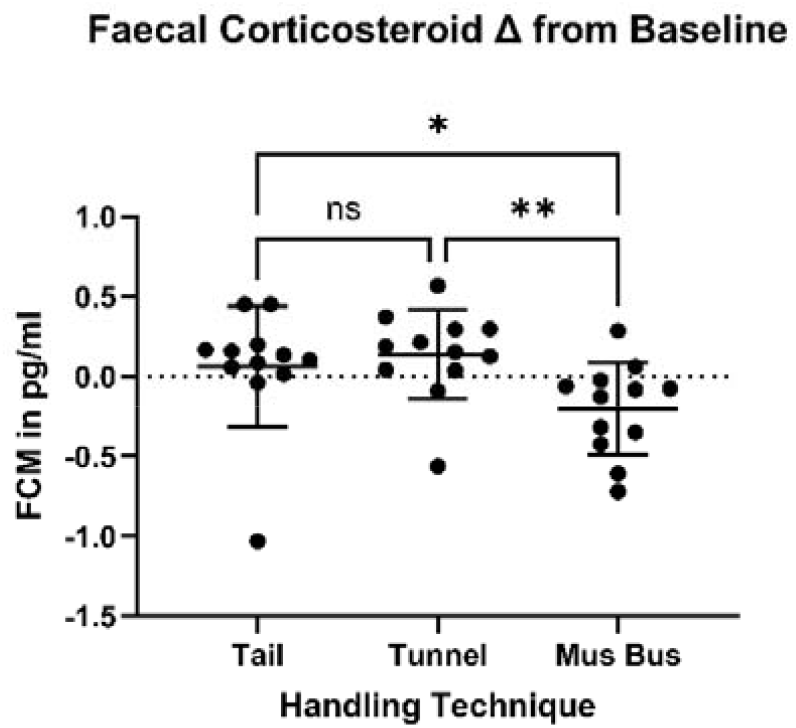
Fecal Corticosteroid Metabolites (FCM) changes from the average baseline values was significantly different between groups. The Mus Mobile group had significantly lower FCM when compared with the other two groups. (p< 0.0057) (Kruskal Wallis statistic = 10. 33).

### Open field Test

#### General Finding

It was understood that despite the provision of tunnel and Mus-Mobile, the animals preferred to nest on bedding material during resting time. Night videos weren’t obtained for this study to explore the playfulness and total usage during active period of mice.

## Discussion

This is the preliminary study and the first and foremost report introducing a new concept of group handling and a novel technique and device to group handle mice. However, Pregnant animals, pups and other strains in research shall be exposed to group handling to fully understand the utility and effects of the device. Also, it has to be studied how the mice will build nests when nesting pads are supplied in cages where Mus-Mobile is employed. The novel device can be accommodated even in the smallest available mice cage, and moreover, tests have to be done in standard mice cages to evaluate the performance of the device.

Inter individual variability in parameters studied using this device and stress while handling could be a factor when the other labs start to use this technique. Hence, studies also shall be planned to find out about variability induced by these factors in the results. Our study did not compare cupping with the other methods studied. This has also to be carried out in order to fully evaluate the utility and advantages of the device.

The device will increase the complexity of the cage and hence will contribute towards added activity and species-specific behaviour. The effects on bizarre fighting (whether it reduces) and other types of vices/stereotypic behaviors when Mus-Mobile is employed shall also be investigated. Multi centric trials and user surveys has to be expedited to establish the technique of group handling using this device. Reinforcing data on corticosteroid levels is also required.

It has been proven that tail-lifting mice can cause stress, anxiety and aversive behavior, and this finding has made scientists develop less stressful techniques like cupping and tunnel- handling. All these techniques have a disadvantage in that they can only handle mice individually and not in groups. This study produces scientific evidence on the advantages of group handling of laboratory mice for the first time. Group handling using the novel device significantly reduced the time taken and number of attempts taken for cage changes, a major reason that prevented tunnel-handling from being adopted globally. Shifting mice in groups using “Mus-Mobile” resulted in significantly lower fecal corticosteroid levels and higher voluntary interaction time. Open field activity and time spent in the central arena were not altered when compared with tail-lifting and tunnel handling in C57BL/6NCrl mice. Body weights were not different when Mus-Mobile was employed with respect to the other groups. These results are encouraging as it indicates that the functionality in behavioral tests outcomes will not be influenced by group handling. This work shows that group-shifting is the next horizon for animal welfare, as this device can revolutionize the handling of laboratory mice to bring in better animal wellbeing.

## Conclusion

Group shifting of mice can be advantageous in terms of time taken for cage changing and ease of performing cage changings in terms of number of efforts made and this technique results in lesser stress and anxiety. Shifting mice in groups will not affect animal’s body weight or activity levels in open field in comparison with tail handling or tunnel. More studies are required to evaluate the effect of group shifting in pregnant and lactating mice and in young ones.

## Supporting information

Data sets

## Acknowledgments

The authors hereby acknowledge TRC of SCTIMST, Director and Head BMT Wing to fund the work. Our animal handler Shri. Manoj M has been the talent behind the scenes and his commitment is hereby acknowledged. Dr. Lynda V Thomas and Mrs. Rukhiya Salim, Dr Ansar Fasaludeen, Haritha Raj, Anjana VA and Sandra Krishnan are hereby acknowledged for their support to the work.

